# Repertoire-Scale Measures of Antigen Binding

**DOI:** 10.1101/2022.06.24.497473

**Authors:** Rohit Arora, Ramy Arnaout

**Affiliations:** Division of Clinical Pathology, Department of Pathology, Beth Israel Deaconess Medical Center, Boston, MA 02215.; Division of Clinical Informatics, Department of Medicine, Beth Israel Deaconess Medical Center, Boston, MA 02215.; Iktos Inc. 50 Milk Street, Floor 16, Boston, MA, 02109

**Keywords:** B-cell repertoires, T-cell repertoires, Immunological diversity, Gibbs free energy, Antigen binding, Public repertoires, Immunomics

## Abstract

Antibodies and T-cell receptors (TCRs) are the fundamental building blocks of adaptive immunity. Repertoire-scale functionality derives from their epitope-binding properties, just as macroscopic properties like temperature derive from microscopic molecular properties. However, most approaches to repertoire-scale measurement, including sequence diversity and entropy, are not based on antibody or TCR function in this way. Thus, they potentially overlook key features of immunological function. Here we present a framework that describes repertoires in terms of the epitope-binding properties of their constituent antibodies and TCRs, based on analysis of thousands of antibody-antigen and TCR-peptide-major-histocompatibility-complex binding interactions and over 400 high-throughput repertoires. We show that repertoires consist of loose overlapping classes of antibodies and TCRs with similar binding properties. We demonstrate the potential of this framework to distinguish specific responses vs. bystander activation in influenza vaccinees, stratify CMV-infected cohorts, and identify potential immunological “super-agers.” Classes add a new dimension to assessment of immune function.

## Introduction

Repertoires are routinely characterized according to the number and frequency of unique V(D)J-recombined antibody and TCR gene sequences they contain (henceforth “genes;” Fig. 1a). This is known as *sequence diversity,* and is measured using a variety of sequence-based diversity indices, including (species) richness, Shannon entropy (*1, 2*), and others related to Hill’s *^q^D*-number framework (Fig. 1b) (*3*). Sequence-based diversity indices (henceforth “sequence diversity”) have shown promise as biomarkers, for example as predictors of response to cancer immunotherapy (*4*) and as correlates of healthy aging (*5-7*). However, sequence diversity overlooks fundamental features of repertoire function. For example, sequence diversity cannot indicate whether a repertoire with a given number of different genes contains epitope-binding capacity (*8*) for many different epitopes or for only a few (Fig. 1c-d), or how well antibodies or TCRs from a second repertoire might also bind a given set of epitopes (Fig. 1e). The reason for this shortcoming is that sequence diversity measures only the number of different antibodies or TCRs, but not their basic function: epitope binding.

**Figure 1.**
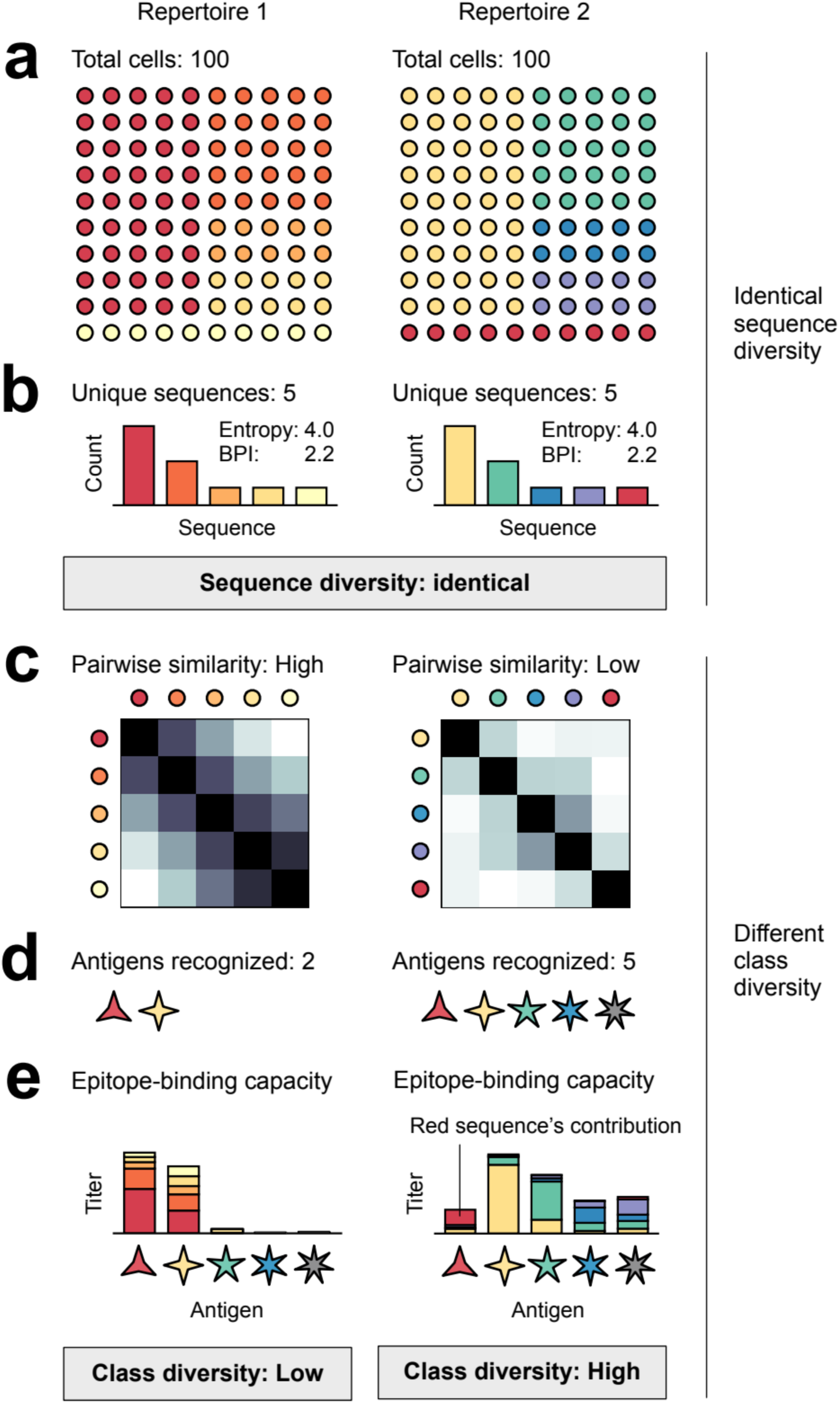
Sequence diversity vs. class diversity Each circle represents a B or T cell; each color represents a unique antibody or TCR sequence. Similar colors encode antibodies or TCRs with similar epitope-binding properties. Two repertoires, e.g. Repertoires 1 and 2 **(a)**, that have the same total number of cells (a) and identical sequence-frequency distributions **(b)**, have identical sequence diversity (for all *^q^D*); insets give the effective-number versions (*3, 31, 32*) of entropy and BPI, ^1^*D*=e*^Shannon^ ^entropy^* and ^∞^*D*=1/BPI. Lower pairwise binding similarities in Repertoire 2 **(c)** give Repertoire 2 higher class diversity than Repertoire 1; Repertoire 2 can recognize more different epitopes **(d)**. Color coding reflects optimal binding (e.g. red sequence, red epitope). The colors of the bars in **(e)** indicate the contributions of the antibody or TCR encoded by the sequence of that color. Similar colors bind better than different colors. Higher frequencies (b) can partially compensate for weaker binding.

Epitope binding—of antibody to antigen or of TCR to peptide-major histocompatibility complex (pMHC)—is routinely measured using dissociation constants (*K_d_*), for example to determine which of several antibodies has the highest affinity for a given epitope (*9, 10*). (Another common measure is IC_50_, used in inhibition experiments.) *K_d_* is related to the Gibbs free energy of binding (*ΔG*) by the equation *ΔG*=-*RT*ln(*K_d_*), where *R* is the gas constant and *T* is the temperature, illustrating the relationship between *K_d_* and thermodynamic first principles. In immunology, it is widely understood that antibodies or TCRs with similar gene sequences often have similar *K_d_* for a given set of antigens or pMHCs (*11-13*), even as targeted substitutions of amino acids can change *K_d_* enough to effectively abolish binding (*14, 15*) (binding is “error-tolerant but attack-prone” (*16*)). Binding similarity among antibodies or TCRs (Fig. 1c) is the basis of phenomena fundamental to adaptive immunity, including polyspecificity/cross-reactivity and degeneracy/redundancy (*17, 18*). These phenomena are what allow so-called natural antibodies (IgM) to recognize many different antigens despite relatively low sequence diversity, with large numbers of antibodies of similar specificity compensating for individually weak *K_d_*S (*19, 20*). Thus, in a qualitative sense, the idea that binding similarities between antibodies or TCRs can, in the aggregate, have important repertoire-scale effects is well established (Fig. 1d-e) (*21*). We sought to develop this idea quantitatively, by developing quantitative repertoire-scale measures based on the binding properties of repertoires’ constituent antibodies and TCRs.

## Materials and Methods

### Overview (Fig. 2a)

391 IGH and TRB repertoires were obtained from 202 human subjects, as previously described (“High-throughput repertoires”). The ratio of dissociation constants was used as the definition of binding similarity (“Definition of binding similarity”), which is a simple log transformation of ΔΔG. Linear and nonlinear models estimating this ratio, using edit distance and/or amino-acid biophysical properties as components, were fit to the experimental *ΔΔG* values in the SKEMPI 2.0 database (“Experimental binding data,” “|*ΔΔG*| distribution” sections below), with a mean model being chosen for further analysis (“Model selection”). (Note the difference between using an input, e.g. edit distance, as the output, *y*=*x*, and using a model fit on data as the output, *y*=*f*(*x*).) *^q^D* and *^q^D_S_* values were calculated as previously described, with *^q^D* values corrected for sampling using the Recon software package (“Diversity measures”). *^q^D_S_* values were tested for robustness to sampling using metarepertoires constructed by pooling individual repertoires and subsampling (“Robustness to sampling”) and, separately, by measuring the extent to which relative *^q^D_S_* of a pair of repertoires is preserved upon sampling (“Robustness of relative ordering of *^q^D_S_* as a function of sample size”). Validity was established using *in silico* repertoires (“*In silico* repertoires”).

**Figure 2.**
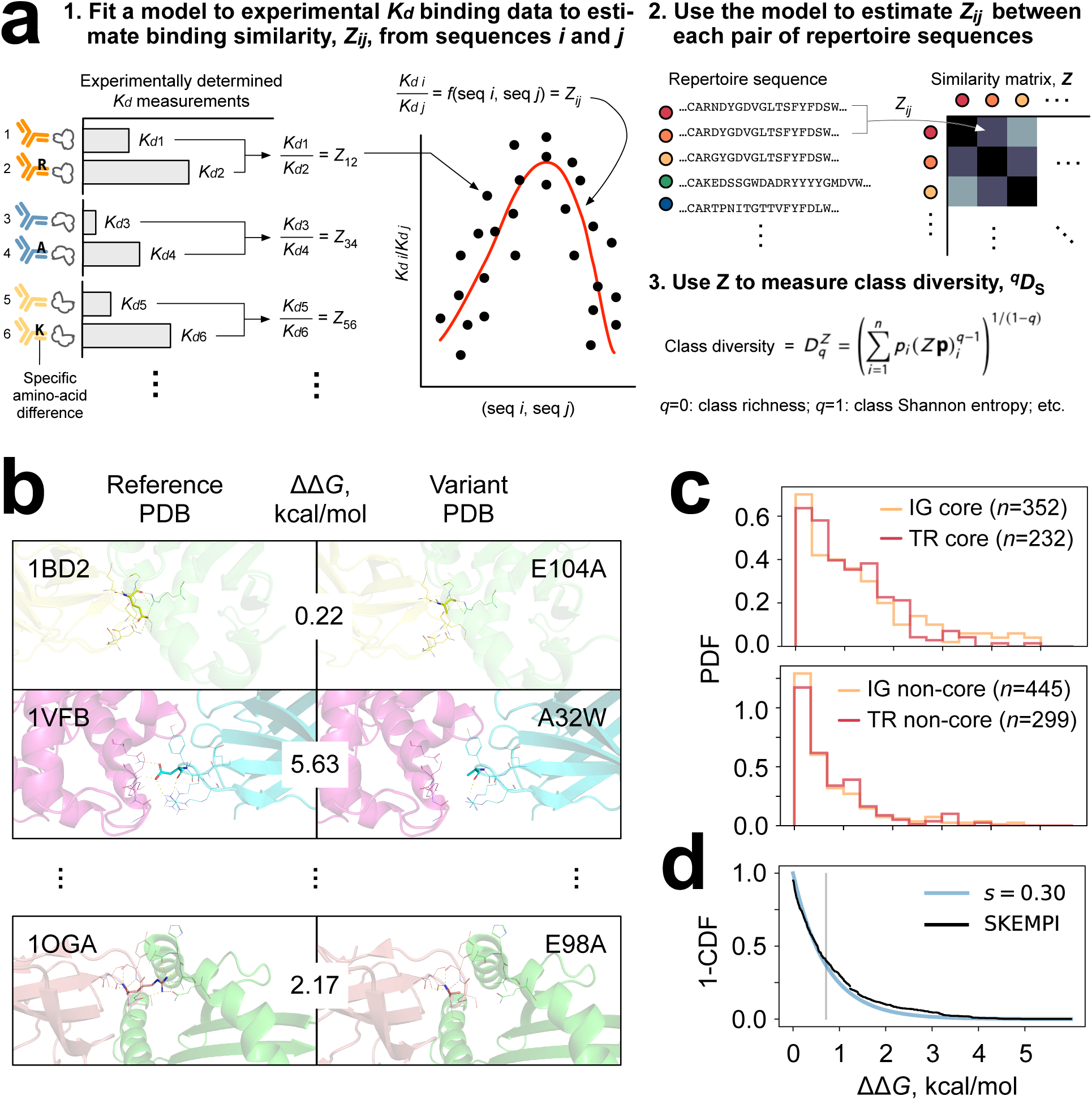
Large-scale experimental *ΔK_d_* for single-amino-acid substitutions on binding **(a)**Overview. In this study, the experimental *K_d_* binding data is from SKEMPI. **(b)** Examples of reference-variant pairs with the view centered on the substituted amino acid. PDB ID in upper left of each row; substitution in upper right. **(c)** Distributions for core (top) and non-core (bottom) mutations for immunoglobulin (IG) and TCR (TR) pairs. **(d)** Combining the distributions in (b) proportional to the relative frequencies of core and non-core residues results in an overall distribution (black), plotted as 1 minus the cumulative distribution function (CDF) and an exponential fit (blue, *e*^1/(-*RT*ln*s*)^). Gray line indicates the mean –*Rt*ln(*s*).

### High-throughput repertoires

391 quantitative high-throughput IGH and TRB repertoires were obtained from 202 human subjects. These included IgH from naïve and memory B cells from DNA (*n*=3 individuals) (*39*); TRB chains from DNA from healthy subjects known to be serologically negative for cytomegalovirus (CMV) (*n*=69 individuals) (*49*) and from healthy subjects whose CMV serostatus was unknown (*n*=41 individuals) (*5*); pooled barcoded IGG and IGM heavy chains from mRNA from healthy subjects before and seven days after administration of one of two influenza vaccines (*n*=28 individuals) (*40*); quantitative pooled TRB chains from DNA for subjects who were otherwise healthy but serologically CMV positive (*n*=51 individuals) (*49*) (a batch processing effect was discovered in which singletons were removed from the other ∼400 repertoires in this dataset, obstructing comparison and limiting us to 69+51=120 repertoires from this dataset); and IGH chains (all isotypes) from DNA for subjects enrolled in the Multi-Ethnic study of Atherosclerosis (MESA; *n*=41 individuals) (*59*). CDR3 annotation was performed using our in-house pipeline as previously reported (*61*) and standard tools (e.g. IMGT (*81*)). Details for obtaining these datasets are available from the references.

### Definition of binding similarity

The ratio of dissociation constants was used as the definition of binding similarity (see Results, “A quantitative definition of binding similarity between two antibodies or TCRs,” for motivation). This ratio is related to ΔΔG by exponentiation: *K_d_*1/*K_d_*2 = e^ΔΔG/RT^. A model estimating this ratio was fit to the experimental *ΔΔG* values in the SKEMPI 2.0 database (*11*) as described below.

### Experimental binding data

Each SKEMPI entry included a Protein Data Bank (PDB) identifier (*27*), the type of structural region (*25*) that contains the substitution(s), one or more PDB coordinates, and in nearly all cases the dissociation constant (*K_d_*) of each member of the pair (referred to in the database and Fig. 2b as “wild type” and “mutant”). The Structural Antibody Database (*82*) and the Structural TCR Database (*83*) were used for assigning species. SKEMPI entries were extracted for all single amino-acid substitutions for which *K_d_* for both wild type and mutant were recorded and |*ΔΔG*| was calculated.

### |ΔΔG| distribution

Only entries that involved binding between antibody and antigen (n=797) or TCR and pMHC (n=531) were considered (total *n*=1,328). Following earlier observations about the heterogeneity of effects of amino-acid substitutions depending on their structural position within the binding interface (“core” vs. “non-core” (*25*)) entries were split into core (*n*=584) and non-core (*n*=744) groups. Distributions for these were confirmed to differ substantially from each other (Mann-Whitney U [MWU] p-value 2.0x10^-33^), with substitution of core residues having a 13-fold (geometric) mean effect on binding (*84*) and non-core residues a 4-fold effect. Both distributions were long-tailed (Fig. 2c) and reasonably well described by exponentials (i.e., equations of the form *ke^-kx^*, with the value of *k* depending on the specific distribution). Distributions for antibody-antigen (*n*=352) and TCR pMHC (*n*=232) core residues were statistically indistinguishable from each other (MWU *p*=0.21), as were distributions for antibody-antigen vs. TCR pMHC non-core residues (n=445 for antibody-antigen and 229 for TCR-pMHC; MWU *p*=0.13). However, core differed from non-core distributions (MWU *p*=1.12×10^-6^-7.37×10^-9^). These results held separately for human and non-human proteins (nearly all of which were from mouse, *Mus musculus*). Detailed manual review of nine structures containing substitutions in human IgH or TCRβ CDR3s (1BD2, 1OGA,473 3BN9, 3QDJ, 3SE8, 3SE9, 4I77, 5C6T, 5E9D) using PyMol v2.2 (*85*) revealed fairly constant proportions of core vs. non-core residues, consistent with the general features of immunolgobulin-receptor superfamily interactions—specifically, 0.15±0.05 of CDR3 amino acids consisted of core residues vs. 0.85±0.05 non-core, with no obvious difference between chain types—and so core and non-core distributions were combined with a weighting of 0.15:0.85. The resulting distribution of |*ΔΔG*| values was again long-tailed and was well fit well by an exponential until ∼3.2 kcal/mol, after which mutations with extreme effects on binding were modestly but clearly overrepresented relative to the exponential model. A review of sources cited by SKEMPI suggested ascertainment bias as the explanation: targeted/selective experimentation on amino acid substitutions with unusually strong effects (e.g. refs in (*86*). To counter this bias, these extreme values (beyond 3.2 kcal/mol) were removed. As a sensitivity analysis, different cutoffs were tested; all reported results were robust to extreme-value cutoffs from 3.0 to 3.4 kcal/mol.

### Model selection

Each of several models was evaluated by bootstrap with random 2:1 training:test-set splits. Each model was fit by minimizing root mean-squared error (RMSE) of |*ΔΔG|* on a random 2/3 of the data (minus extreme values; see previous section) and tested by calculating RMSE on the remaining 1/3. Each fit was repeated 200 times and mean and standard deviation of the fit parameters and RMSE was recorded.

For models based on amino-acid biophysics, after extensive review (*29, 87–89*) the following raw measures from (*28*) were initially included: side-chain molecular weight, side-chain van der Waals volume, NMR measures NM1, NM7, and NM12, side-chain total surface area, polar surface area, polarizability, electronegativity, number of hydrogen-bond donors, number of hydrogen-bond acceptors, number of positive charges, and number of negative charges. The latter two were combined into a single “charge” variable as no. positive minus no. negative (no information is lost in this process, since none of the 20 standard residues has both positive and negative charges). Because a large number of hydrophobicity scales exist, these were systematically analyzed instead of simply also including TL (for thin-layer chromatography) and logP from (*28*). Based on over 100 such scales examined in (*90*) (Table V therein), the seven least-mutually-redundant scales were selected for inclusion (*91-97*) (Tables IV, III, I, 2, 3, 2, and 1 of these references, respectively), resulting in 19 measurements for each of the 20 canonical amino acids. All-pair correlations revealed several pairs of measurements had *R*^2^≥0.85. A single member of each such pair was retained, resulting in 14 fairly independent (median/interquartile range for pairwise *R*^2^=0.09/0.32) measures (“properties”). PCA was performed and the first five principal components were used (“PCs”; variance explained, 91%, comparable to the first five PCs in (*30*)). Linear fits were performed on properties (RMSE 0.72±0.04 kcal/mol) and on PCs (RMSE 0.72±0.05 kcal/mol), both using ordinary least squares. Nonlinear models were fit using support-vector regression with RBF kernel trained on either properties (0.79±0.05 kcal/mol) or PCs (RMSE 0.78±0.06 kcal/mol). the performance of models that were trained on fewer properties or fewer PCs (e.g. molecular weight, polar surface area, and electronegativity, the major contributors to the first three PCs) was statistically indistinguishable from the above.

For the mean model, the fit was the mean |*ΔΔG*| of the training set (RMSE, 0.71±0.03 kcal/mol), corresponding to *K_d_*1/*K_d_*2=*s*=0.30 (95% CI 0.28-0.32). Low RMSE and simplicity favored the mean model, and so it was used. To assess how this single-substitution model generalized to multiple substitutions, the limited multi-substitution SKEMPI entries were tested and found compatible with a multiplicatively independent model, *Z_ij_*,=*s^m^*, with *Z_ij_*, the similarity between antibodies or TCRs *i* and *j* and *m* the number of amino-acid differences between them.

### Diversity measures

*^q^D* was calculated as previously described (*37*) and *^q^D_S_* was calculated according to Leinster and Cobbold (*31*) following Eq. 1. *^q^D* was corrected for sampling error using Recon v3.0 (github.com/ArnaoutLab/Recon; default settings) as previously described (*37*). For readability, the notation was changed from *^q^DZ* in (*31*) to *^q^D_S_* and from ***Zp****i* to ***S****i* (*S* for “similarity”). Histograms confirmed that the vast majority of off-diagonals were always close to zero, allowing sensitivity to *q* (*98*). Note Hill’s framework (*3*) has inspired several methods for incorporating similarity into diversity measurements that retain useful features of Hill’s framework (*74*). Two such frameworks were introduced with explicit discussion of how to decompose population-level diversity into within- and between-group components (*98, 99*). Each has advantages. Leinster and Cobbold’s was chosen here for ease of applicability and interpretability.

### Robustness to sampling (Fig. 3a-c)

IGH and TRB were analyzed separately. A conservative upper bound for IGH was evaluated by constructing a metarepertoire by combining the following: IGG sequences of subjects before vaccination (n=28 individuals) (*40*), sequences from memory cells from healthy subjects from public database (n=3 individuals) (*39*), and sequences from subjects enrolled in MESA study (n=41 individuals) (*59*). Sequences were sampled as singletons from this set, since repertoires of all-singletons will have higher diversity than repertoires with larger clone sizes. TRB metarepertoires were constructed by combining sequences from CMV seronegative individuals (n=69 individuals) (*49*) and again sampling at uniform frequency. CMV seronegative individuals were preferred for their higher diversity. Samples from real-world repertoires were from subject D3 for IgH (from DNA), subject SRR960344 for IGH (from mRNA), and subject Keck0070 for TRB (CMV seronegative) from the references above. For these samples, genes were sampled proportional to their frequency in the repertoire. The results from this analysis are conservative because they assume that a given person’s repertoire is as diverse as the combined repertoires of the 72 (IGH) or 69 (TRB) repertoires above; in reality, no single person’s repertoire is likely to be this diverse, meaning that sampling from a single person’s repertoire will be more robust than the results of this analysis.

**Figure 3.**
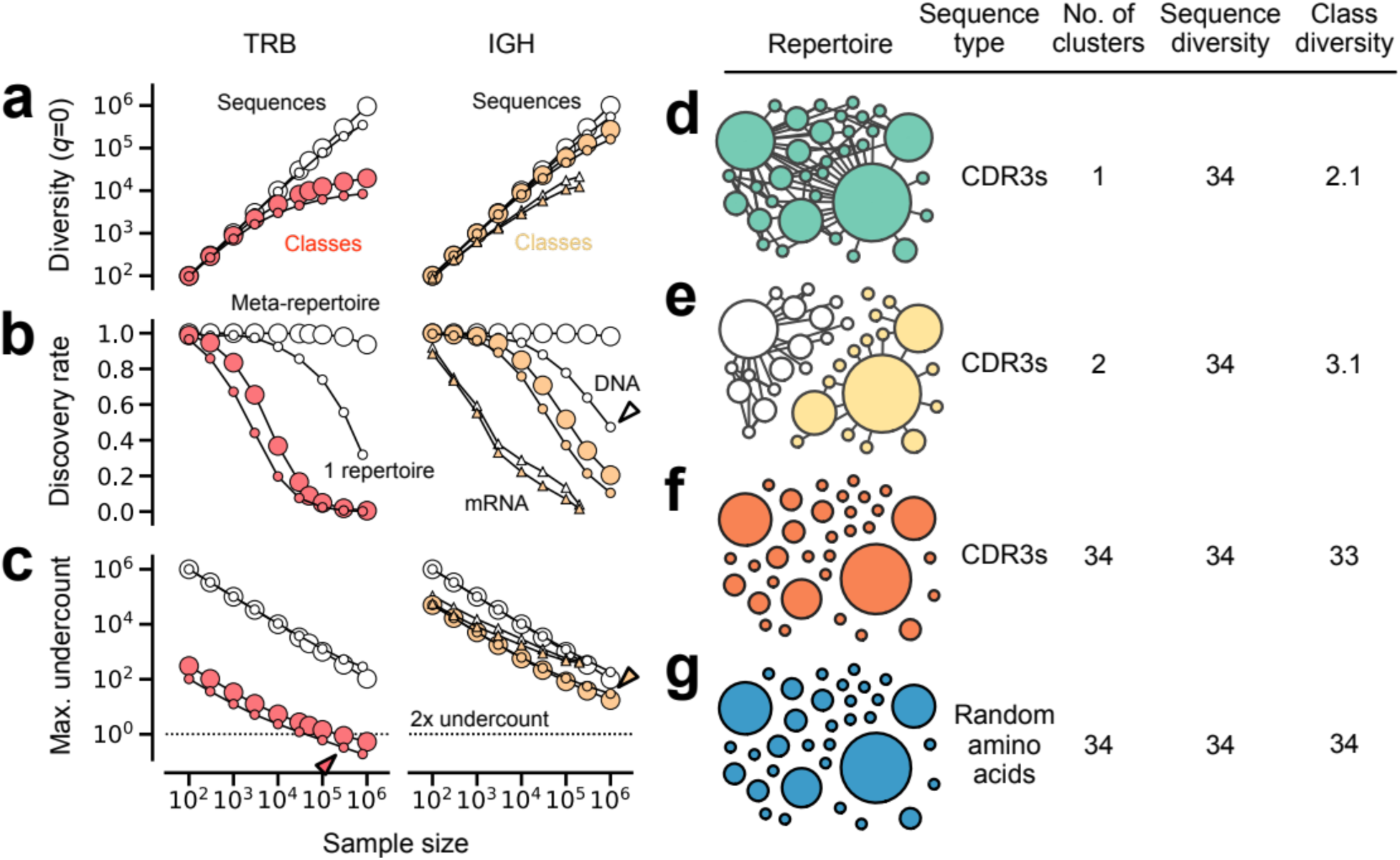
Robustness and validity **(a)** ^0^*D* and ^0^*D_S_* diversity, **(b)** discovery rate, and **(c)** maximum error for sequences (open symbols) and classes (filled circles) for repertoires from DNA (small circles) or mRNA (small triangles) and for meta-repertoires (large circles) vs. sample size. Maximum undercount in (c) is the maximum fraction by which sample diversity will underestimate overall diversity (*37*). Red arrowhead: underestimate for a 300,000-sequence TRB repertoire is ≤33%. Yellow arrowhead: sample class diversity of a 1-million-sequence IGH repertoire will underestimate overall class diversity by ≤30x. Open arrowhead: for a million-sequence IGH repertoire from DNA, there is a ∼50-50 chance that the next sequence will be new. **(d)**-**(f)**, validity: sequence vs. class diversity for four *in silico* repertoires, each with 34 unique/752 total sequences with identical sequence-frequency distributions (c.f. Fig. 1b). In the networks, each node represents a unique sequence; node size reflects that sequence’s frequency in the repertoire. Edges connect sequences that differ at a single amino-acid position. **(d)** CDR3s from a somatically hypermutated IGG clonotype. The extent to which class diversity exceeds 1 reflects intra-clone diversity. **(e)** CDR3s from two different IGG clonotypes. **(f)** CDR3s drawn randomly from repertoires in this study. **(g)** Non-CDR3 amino acid sequences generated uniformly at random.

### In silico repertoires (Fig. 3d-g)

Small synthetic *in silico* repertoires were created by sampling from post-influenza-vaccination IgG repertoires (*40*). A single clone was chosen at random to serve as a reference repertoire (34 unique/752 total CDR3s; CDR3 length, 17 amino acids). A network representation was created in which the nodes represent unique CDR3s and each pair of nodes of edit distance 1 is connected by an edge. To create a two-clone repertoire with identical node-size distribution and intra-clone edges as the reference repertoire, half of the unique CDR3s of the reference repertoire were replaced with CDR3s from a second, unrelated clone chosen at random from another repertoire. CDR3s from the second clone were filtered to induce a subgraph identical to the subgraph of the replaced CDR3s. Frequencies of the replaced CDR3s were assigned to the new CDR3s. A third repertoire comprising unrelated CDR3s was created by sampling 34 length-17 CDR3s at random from among all antibody repertoires used in this study (we confirmed that all pairs had edit distance >1, as expected from such sampling), and again assigning these the same frequencies as in the reference repertoire. A fourth repertoire was created from random 17-mer amino-acid sequences, assigning frequencies as above.

### Robustness of relative ordering of *^q^D_S_* as a function of sample size

For a given *q*, for all pairs *i*, *j* of the six large naive and memory B-cell repertoires in (*39*), the ratio *^q^D_Si_*/*^q^D_Sj_* was calculated, indicating which repertoire was more diverse and by what factor (the ground truth or “correct relationship;” e.g. a ratio of 1.3 meant repertoire 1 was 30% more diverse than repertoire 2). For decreasing sample sizes (300,000, 100,000, 30,000, 10,000, 3,000, 1,000, 300, and 100 sequences), each repetoire was randomly subsampled 20 times, *^q^D_S_* was calculated for each subsample, and *^q^D_Si_*/*^q^D_Sj_* was calculated on each pair *i*, *j* of subsamples. For each sample size, the fraction of comparisons that gave the correct relationship was recorded, as well as the mean and standard deviation of the ratios. This showed that for ^0^*D_S_*, differences of ≥50% can be detected at sample sizes of 3,000 sequences, 10% at 30,000 sequences, and 1% at 100,000 sequences, all with 99% confidence (and similarly for ^1^*D_S_*). The same procedure was carried out for TCR using the six largest repertoires from among the 69 obtained from (*49*) from (Keck 069, 070, 080, 093, 095, and 113; maximum sizes of 300,000-500,000 sequences), demonstrating that differences of 3% (^0^*D_S_*) and 4% (^1^*D_S_*) can be detected reliably at sample sizes of ≥50,000.

### CMV classifier (Figs. 4d, 4f-g)

A random-forest classifier was trained (using scikit-learn’s RandomForestClassifier module) to predict the probability that a repertoire was CMV seronegative. The model was fit on *^1^D_S_* (class Shannon entropy), ^∞^*D* (a measure of how large the largest *clone* is), and ^∞^*DS* (a measure of how large the largest *class* is) with a training:test set split of 2:1, 30 estimators, and a maximum depth of 1. Classifier AUC was 0.83. Other parameters were tested with indistinguishable results (e.g. using ^0^*D_S_* instead of ^1^*DS*; using a maximum depth of 2 instead of 1). 90% confidence thresholds for CMV positivity and negativity were used to annotate repertoires in Figs. 4f-g (plus and minus signs). If a repertoire did not meet that threshold, it was left unannotated (no sign).

**Figure 4.**
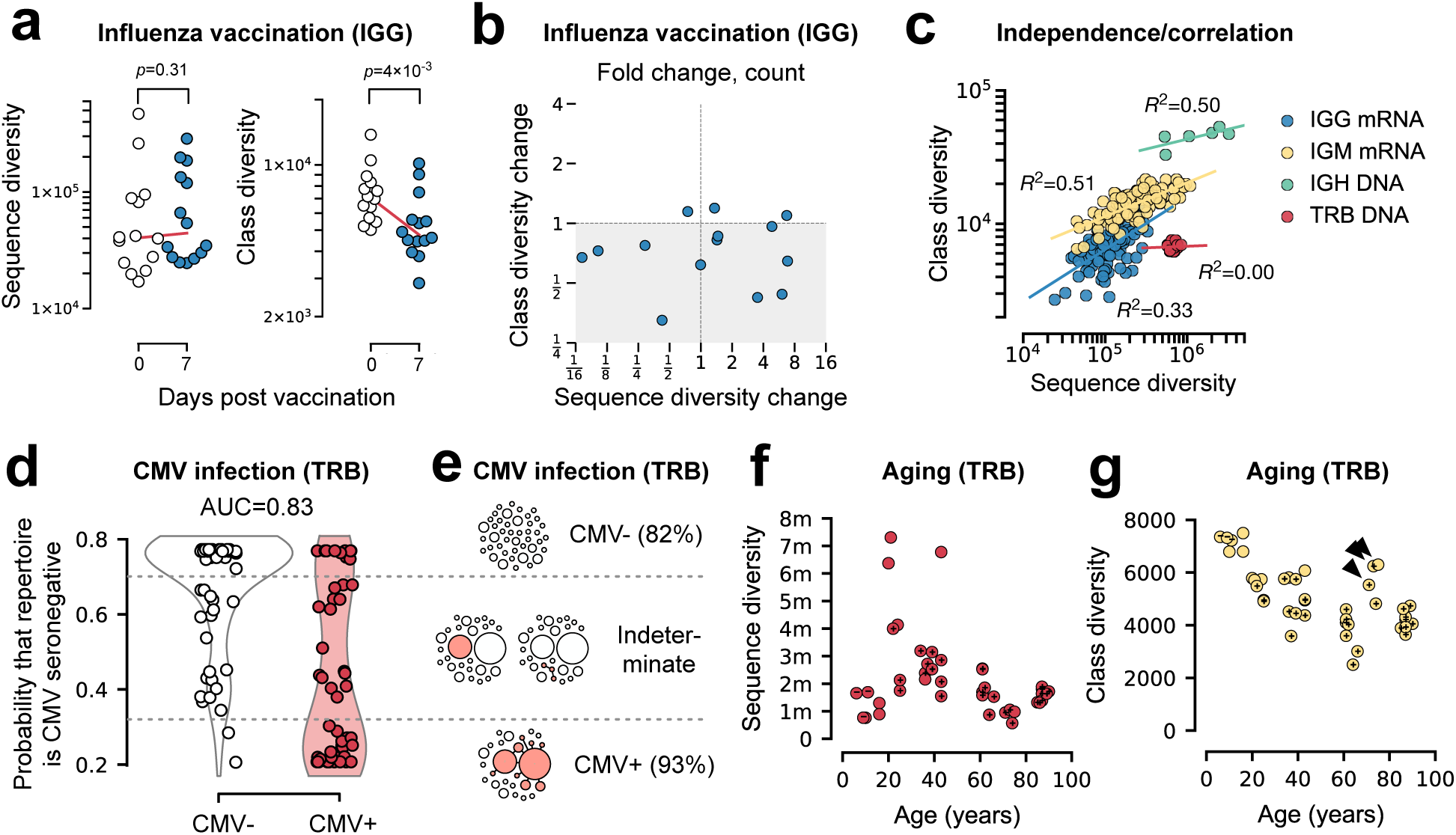
Class diversity for stratification and discovery **(a)** Sequence and class diversity for IGG repertoires from mRNA in influenza vaccination. **(b)** Fold change in sequence diversity vs. class diversity. **(c)** Class diversity vs. sequence diversity for different repertoire types. Each symbol is a repertoire. **(d)** Score from a random-forest classifier based on combining sequence and class diversity for TRB repertoires from DNA for *q*=1 and ∞ for CMV-seronegative (empty circles) vs. seropositive (filled circles) individuals, with **(e)** network-diagram schematics of the corresponding repertoires; filled circles represent CMV-specific CDR3s. Note the top row of CMV-repertoires in (d) contains 51 repertoires (34 overlapping symbols). **(f)** Sequence diversity and **(g)** class diversity for TRB repertoires from DNA by age. Arrowheads indicate exceptional individuals. Dotted lines, linear and exponential fits; + and – designate 90%-confidence imputed CMV serostatus. *q*=0 except in (d).

### Scientific software

Recon v3.0 was performed using Python 3.7.6 with NumPy version 1.18.0 and SciPy version 1.4.1. All other analyses were performed on Python 3.9.1 with NumPy 1.20.0 and SciPy 1.6.1.

## Results

### Overall approach (Fig. 2a)

We started from the principle that antibodies or TCRs with binding properties similar to those already present in a repertoire should contribute less to the overall diversity of the repertoire than antibodies or TCRs with different binding properties. We developed this principle into repertoire-wide measures in three steps. The first was to derive a quantitative definition of the binding similarity between any two antibodies or TCRs. The second was to develop a method for estimating this quantity for every pair of antibodies or TCRs in a repertoire. The third was to sum contributions to the overall diversity, weighting each antibody or TCR according to the uniqueness it adds, such that antibodies or TCRs that are similar to each other contribute less and those that are different contribute more. Sums of this kind constitute the desired family of repertoire-scale binding measures.

### A quantitative definition of binding similarity between two antibodies or TCRs

There are many sensible ways to define similarity between two antibodies or TCRs. Many are expected to correlate with antigen binding, and indeed some have been shown to do so (*12, 13*). We sought to derive a definition from thermodynamic first principles. We observed that, for two antibodies present at the same concentration, if one antibody or TCR binds its target *n* times better than another, the second antibody or TCR will bind 1/*n*^th^ of the target (*22*). For example, two antibodies that each bind a target half as well as a third antibody collectively have the same binding capacity as that third antibody. Quantitatively, the definition of binding similarity that has this additive property is the ratio *K_d_*1/*K_d_*2, where *K_d_*1 is the dissociation constant of one antibody or TCR for a target and *Kd2* is the dissociation constant of the other antibody or TCR for that target, with *K_d_*1 the smaller of the two. (For completeness, this formulation can be extended across all possible targets; for convenience, because similar antibodies or TCRs usually show similar binding patterns across targets, we treated the ratio for a single target as representative of the relationship, and leave formal extension for future study.) Note that the ratio *K_d_*1/*K_d_*2 is related to the absolute difference in free energy for the two binding interactions, |*ΔΔG|*, by the equation ΔΔ*G*=−*RT*ln(*K_d_*1/*K_d_*2), where *T* is standard temperature (298K) and *R* is the ideal gas constant (1.99×10^-3^ kcal·mol-1·K^-1^).

### A model for estimating binding similarity between pairs of antibodies or TCRs

To estimate *K_d_*1/*K_d_*2 for a given pair of antibodies or TCRs, we fit a model to a large set of experimentally determined measurements of *ΔΔG* for a pair of antibodies or TCRs and a specific binding target. Because *K_d_*1/*K_d_*2 cannot yet be predicted precisely for a given reference-variant pair (*23, 24*), the model’s estimates were expected to be imprecise for any given pair of antibodies or TCRs; however, the law of large numbers—on the order of 10^10^ pairwise comparisons per repertoire—provides for accuracy of the desired overall measures.

The experimental measurements used to train our model consisted of 1,328 systematic comparisons between pairs of *K_d_*S (*11*) measured as ΔΔ*G=*−*RT*ln(*K_d_*1/*K_d_*2). Here, *K_d_*1 is the *K_d_* for the interaction between a given antigen or pMHC epitope and a reference antibody or TCR, *K_d_*2 is the *K_d_* for the interaction between the same epitope and a variant antibody or TCR that differs from the reference by a single amino-acid substitution. Because, on average, amino acids in the interior of protein-interaction interfaces (“core”) are known to affect ΔΔ*G* less than those at the surface (“non-core”) (*25, 26*), we evaluated core and non-core substitutions separately. As expected, we found ΔΔ*G* differed substantially between core and non-core substitutions, both for human antibodies (median/interquartile range 1.05/1.36 kcal·mol^−1^ for *n*=154 core amino-acid substitutions vs. 0.43/0.92 kcal·mol^−1^ for *n*=244 non-core substitutions; Mann-Whitney U *p*-value=3×10^-8^) as well as for human TCRs (0.93/1.22 kcal·mol^−1^, *n*=217 vs. 0.53/0.95 kcal·mol^−1^, *n*=242; MWU *p*=1×10^-6^). Consistent with antibodies’ and TCRs’ structural similarities as members of the immunoglobulin superfamily, we also found that for each subset, ΔΔ*G* distributions were statistically indistinguishable between antibodies and TCRs (*p*=0.21 for core and 0.13 for noncore substitutions by Mann-Whitney U), allowing pooling across IG/TCR for greater statistical confidence. Manual review of crystal structures (*27*) showed that core residues comprised 15±5% of the third complementarity-determining regions (CDR3s) of human immunoglobulin heavy chain (IGH) and TCR-β chain (TRB). Thus, a master distribution of effects of single amino-acid substitutions in IGH or TRB CDR3s (Fig. 2d) was created as a 0.15:0.85 weighted sum of the observed effect sizes for core and non-core substitutions.

We evaluated several multi-parameter statistical models based on specific amino-acid substitutions, including a linear-regression model based on biophysical properties of the substituted amino acids (e.g. molecular weight, electronegativity, and 13 others (*28, 29*); RMSE, 0.70±0.04 kcal·mol^-1^), a linear-regression model based on PCA-dimensionality-reduced aggregate biophysical descriptors (*30*) (5 parameters; RMSE 0.70±0.03 kcal·mol^-1^), and corresponding nonlinear models (RMSE 0.79-0.80±0.05 kcal·mol^-1^). However, in cross-validation none of these models statistically outperformed the simplest possible model, a simple mean (0.71±0.02 kcal·mol^-1^). Therefore, this latter was used. The average similarity for a single amino-acid substitution *s*=*K_d_*1/*K_d_*2=0.30 (95% CI, 0.28-0.32). Comparison to cases in which reference and variant antibodies or TCRs differed by multiple substitutions supported a multiplicative model for the pairwise similarity between IGH or TRB CDR3s that differ at multiple positions. Thus in our model the similarity between two IGH or TRB *i* and *j*, *Z_ij_*, is the average *K_d_*1/*K_d_*2 for an amino acid substitution, *s*, raised to the edit distance, *m*: *Z_ij_*=*s^m^*, with *c*=0.30 based on large-scale experimental binding data, and with that data currently insufficient to justify further model complexity. Results were robust to sensitivity analysis.

### Class diversity from pairwise similarity measures

To obtain the desired repertoire-scale measures for antibodies or TCRs, a sum is taken over all unique pairs of antibodies or TCRs. This sum yields the *effective number* (*3, 31, 32*) of different antibodies or TCRs in the repertoire, taking similarities into account (see Fig. 1c-d). (The effective number is the same as *^q^D_S_* below.) The effective number can be understood in several ways: as a measure of how much of antigen space a repertoire can address; as the number of clusters in the repertoire, discounting overlap between clusters; or the number of completely unrelated antibodies or TCRs a repertoire could be replaced by (antibodies or TCRs with disjoint or completely non-overlapping binding specificities), and still bind the same targets at the same aggregate strength. For example, if an antibody repertoire has 50,000 unique sequences but these all bind the same 2 structurally completely unrelated antigens, the repertoire’s effective number is 2; such a repertoire could be replaced by another repertoire with 2 completely unrelated antibodies, one that binds each antigen.

Pairwise similarities *Z_ij_* for all antibodies or TCRs in a repertoire were calculated using the model, and then summed according to Eq. 1 (*31*):

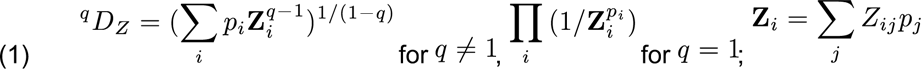

Here, *p_i_* is the frequency of the *i*^th^ antibody or TCR in the repertoire and *q* is the so-called viewpoint parameter, which up-weights antibodies or TCRs based on frequency, just as it does in the Hill framework (where *q*=0 corresponds to species richness; *q*=1, Shannon entropy; *q*=2, the Gini-Simpson index; and so on up to *q*=∞ for the Berger-Parker index (*32*); see e.g. Fig. S1). Setting *q>*0 up-weights higher-frequency antibodies or TCRs to focus on e.g. larger B- or T-cell clones or higher-titer antibodies. *q*=0 yields the unweighted sum. The formulation and notation of *^q^D_S_* were developed in ecology as extensions of the Hill framework (*31*); *^q^D_S_* reduces to *^q^D* if binding similarity is ignored, which is accomplished mathematically by setting ***Z***=***I***. Usefully, this framework is modular: other models of similarity can be explored by simply changing the values of ***Z***.

The new *^q^D_S_* measures estimate the *class diversity* of a repertoire. A class is a set of antibodies or TCRs that have a similar binding pattern (Fig. 1c). Class members bind the same antigens or pMHCs similarly well. Like binding, class membership is not binary but continuous: two antibodies or TCRs are members of the same class to the extent that their binding patterns are similar to each other. A class is an example of what is known in mathematics as a *fuzzy set* (*33*). In immunology, this concept has long been viewed as an organizing principle of antibody and TCR repertoire composition (*34-36*). Our class diversity framework develops this concept into a family of quantitative repertoire-scale measures that can be easily applied, compared, and interpreted in biological and clinical settings, for example for stratification of patient cohorts, as we demonstrate below. We found that *^q^D_S_* is robust to sampling error for sample sizes ≥50,000 T or 100,000 B cells, unlike *^q^D*, which is much more prone to sampling error and is dangerous to use without correction (Fig. 3a-c) (*37, 38*). We also found that the relative order of diversity values—whether repertoire 1 is more diverse than repertoire 2, for a given pair—is preserved in subsamples, such that a 1% difference in BCR and a 5% difference in TCR are detectable at sample sizes of 100,000 and 50,000 sequences, respectively, at 99% confidence, for ^0^*D_S_* and ^1^*DS*, meaning that statements about which of two repertoires is more diverse can be made quite precisely, even when the two values being compared have far higher uncertainty (see Methods). Fig. 3d-g illustrate the relationship between sequence and class diversity (see Methods),

### Class diversity of naive vs. memory B cells

We compared 72 high-throughput CDR3_H_ repertoires from 31 healthy individuals, including three exceptionally deeply sequenced naive (IgM+IgD+CD27-) and memory repertoires from DNA (*39*) and 28 IGM and IGG repertoires from mRNA (Fig. S1) (*40*). (Note that repertoires from mRNA will overrepresent highly transcribing cells such as circulating plasmablasts.) Naive B cells express IGM antibodies, which collectively can bind many different antigens but individually are often weak, polyspecific, and/or degenerate binders (*41-43*). (Note that while up to half of memory B cells express IgM, because naïve cells outnumber memory cells, a majority (∼80%) of IgM+ B cells are naive.) Binding specific antigen may trigger a naive cell to become a memory cell and class-switch to IGG; IGG antibodies are generally stronger binders due to somatic hypermutation and selection, an evolutionary process that diversifies memory lineages away from both the naive repertoire and from each other (*44, 45*). Accordingly, naive repertoires can be considered very diverse in terms of the number of different antibodies they contain but less diverse from a functional perspective to the extent that these antibodies exhibit degeneracy, whereas memory repertoires are very diverse for the distances the evolution of their lineages may have carried them but less diverse insofar as they generally include fewer unique genes (i.e., unique sequences recovered). We found that class diversity reflects these relationships (Fig. S1a,c). For example, we found that even as naive repertoires have 3-10 times as many unique genes as memory repertoires (Fig. S1b,d), memory have nearly as many classes. Thus, while sequence and class diversity are complementary, class diversity better captures the biology and intuition of what it means to be functionally “diverse,” as desired (*46*).

### IGG repertoires in influenza vaccination

To test the potential utility of class diversity as a biomarker, we measured sequence and class diversity on 30 IGG CDR3 repertoires from 14 individuals taken before and seven days after influenza vaccination (Fig. 4a-b) (*40*). In previously exposed populations, vaccination leads to an IGG recall response marked by a rise in sequence diversity at day 7 post administration, as measured by the sequence-diversity index species richness (although these measurements of diversity were not corrected for sampling error in the referenced work) (*40*). However, a rise can result either from clonal diversification, considered a correlate of protection (*47*), or from bystander proliferation (*48*). We hypothesized that class diversity might distinguish between these possibilities: if a rise in sequence diversity is from clonal diversification, class diversity should fall, since the new sequences will be similar to each other; if from bystander effects, meaning unrelated sequences, class diversity should rise. After correcting for sampling error (*37*) we found that sequence diversity rose in only about two-thirds of subjects, likely reflecting a rise in plasmablasts (Fig. 4a-b). However, almost all individuals experienced a fall in class diversity—in many cases, by over a third—and only rarely a rise (never more than 20%; bystander proliferation), suggesting clonal diversification in some but not all vaccinees. Thus, class diversity stratified individuals independently of sequence diversity, which may suggest a marker of successful vs. unsuccessful vaccination.

### TRB repertoires in CMV exposure

Interestingly, this ability to stratify came despite a fairly strong overall correlation between sequence and class diversity for IGG repertoires obtained from mRNA (*R*^2^=0.33) (Fig. 4c). Class diversity also correlated fairly strongly with sequence diversity for IGM repertoires from mRNA (*R*^2^=0.51) and IGH from DNA (*R*^2^=0.50), indicating that sequence diversity explains about half the variance in class diversity in antibody repertoires. However, in TRB repertoires (from DNA) (*49*), class diversity was independent of sequence diversity (*R*^2^=0.00) (Fig. 4c).

We therefore further tested class diversity’s potential for patient stratification by comparing sequence diversity and class diversity in the setting of human cytomegalovirus (CMV) exposure, using TRB CDR3 repertoires from 51 known-seropositive cases and 69 seronegative controls (Fig. 4d-e; see Methods, including note on batch effect) (*49*). CMV is a highly prevalent chronic human herpesvirus infection that can cause life-threatening illness in newborns and transplant recipients and is thought to contribute to heart disease (*50*). The hallmark of CMV exposure is low TRB CDR3 sequence diversity and large T-cell clones, leading to high-frequency CDR3s (*51*). We focused on this feature by considering diversity as measured using the maximum *q*(^∞^*D*; =BPI^-1^) (*52*): the bigger the largest clone, the higher the BPI, and the lower the ^∞^*D*. The class-diversity analog of BPI is class BPI (^∞^*DS*^-1^): high class BPI means a repertoire contains large sets of *similar* TRB CDR3s, regardless of the size of any one clone. (We found that in CMV-seropositive individuals, both ^∞^*D* and ^∞^*DS* trended lower than in controls: thus, in CMV, not only are clones larger, but summed over the repertoire, their CDR3 sequences are more similar than in negative controls (Fig. 4e), despite TCR not undergoing somatic hypermutation as antibodies do. Accuracy was 5% better than in an otherwise identical model that used ^1^*D* and ^∞^*D* but no class diversity measures. Finally, we found that combining these measures with class entropy or class richness improved stratification of CMV status, again demonstrating the potential of class diversity to contribute to diagnostic stratification (Fig. 4d-e).

### TRB repertoires in aging

The potential ability to stratify populations raises the possibility of identifying immunologically exceptional individuals. We tested the potential of class diversity to identify such individuals by measuring sequence and class diversity as functions of age, using TRB CDR3 repertoires from 41 healthy 6- to 90-year-olds (*5*). Before accounting for sampling error, sequence diversity correlates negatively with age, as the thymus involutes and as larger clones displace smaller ones (*5*). After accounting for sampling error (*37*) we found this trend begins only in the fourth decade of life in this dataset, with low values in adolescents and several apparent outliers among 20-40-year-olds (Fig. 4f). In contrast, class diversity was characterized by a steep drop during adolescence followed by plateauing, indicating a relatively rapid loss of functional diversity during this period, followed by relative functional stability even as sequence diversity continue to be lost (Fig. 4g). However, three subjects in their 70s appeared to buck this trend, with sequence richness similar to that of other seniors but with the class richness of children (Fig. 4g, arrowheads). Using Figs. 4d-e, CMV status is unlikely as an explanation. Additional clinical data was unavailable. It is unclear whether the unusually high class diversity of these individuals, who comprise a quarter of individuals ≥65 years of age in this cohort, reflects a transient rise in class diversity or persistence since childhood possibly reminiscent of "super-aging" (*53*). This utility for identifying unusual or exceptional individuals may be useful for revealing heterogeneity in other cohorts as well.

### Implications for public vs. private repertoires

Finally, we found that class diversity also suggests a resolution to the paradox of how it is that, clinically, most people respond with similar success to a given immunological challenge despite sharing few antibody and TCR genes (<1% for IGH (*54, 55*) and <10% for TRB (*56*) CDR3s) and with even lower percent overlap (as shared genes are often low in frequency): the 90-99% of genes that are “private” (*57, 58*) may simply belong to common classes. As a first test of this hypothesis, we generated rarefaction curves for genes and classes using 71 IGH and (separately) 69 TRB repertoires pooled across the population (*5, 39, 40, 49, 59*). We found that the number of genes grew linearly with continued sampling, as expected for low sequence overlap (Fig. 3a, open symbols): most genes were new to the sample, and so the discovery rate remained high (Fig. 3b, open symbols). In contrast, we found that the number of classes saturated for TRB and begun to plateau, with discovery rate of ≤10% at sample sizes of 0.5-1 million cells (Fig. 3b, small symbols), consistent with a very high degree of class overlap between individuals (Fig. 3a-c, filled symbols). Extraordinarily, for both IGH and TRB, the number of classes in the entire sampled population was not substantially larger than that in a single young, healthy individual. This result implies that the private repertoire is functionally public and that classes hold additional useful patterns, which future studies may reveal (*18, 60–62*).

## Discussion

Ultimately, repertoires owe their large-scale organization to patterns of similarity among their constituent antibodies and TCRs. Consequently, similarity has long been of interest. It is the basis of well known, powerful, and insightful coarse-graining techniques such as binning by segment use (*63, 64*), clone collapsing by Hamming distance (*40*) (also used for network clustering (*65, 66*)), and defining amino-acid motifs (*12, 13*). That the results are often specificity groups that contain antibodies or TCRs with functional similarity should not be surprising, since these techniques are all based on sequence similarity (whether in the form of edit distance or a more complex function of e.g. mutation frequencies or biophysical properties), and proteins with similar sequence often have similar properties. Thus all of these techniques can be seen as indirect measures of the same, more fundamental property: similarity of antigen binding.

Quantitatively and thermodynamically, there appears to be only one way of defining antigen-binding similarity between two antibodies or two TCRs that respects the basic property of addition, which is as the ratio of their *K_d_*S for a given antigen (although see limitations below). (A reasonable lower limit on similarity, below which similarity is 0, would be set by antibody solubility; the upper limit is of course 1.) Using this definition, summing yields the total number of unique binders in the repertoire (*31*). Other definitions of similarity can be used to populate the similarity matrix ***Z*** in Eq. (1), but we have reasoned that only the ratio of *K_d_*S yields this straightforward repertoire-scale interpretation.

Note this approach does not restrict the investigator to a specific Hill-type (*3*) diversity measure such as richness, Shannon entropy, or Simpson’s index (*12*). Just as e.g. sequence richness and sequence entropy capture different aspects of a repertoire’s sequence-frequency distribution, and can be useful in different circumstances as a result (*67*), the approach described here allows investigation of whatever aspect of *class*-frequency distribution might be of interest (selectively up-counting larger classes by chosing a larger value of *q*; for example, *q*=1 for ^1^*DS*, the effective number equivalent of class Shannon entropy). The same effective-number interpretation applies (*3, 31, 32*). usefully, note also that the ratio-of-*K_d_*S definition is continuous, meaning that low-level binding similarity among e.g. natural antibodies is not ignored for being below some arbitrary cutoff, as it is in several of the referenced prior techniques; the collective impact of many low-similarity antibodies, for example, is less likely to be overlooked in measurements using the framework we describe. Nature offers several examples of the importance of such “weak ties” (*68, 69*); given the extraordinary number of different genes in immune repertoires, low-level similarities may well add up.

One limitation of this study is that the ratio of *K_d_*S is for a single reference antigen. To be clear, the experimental data on which our model is based includes a very wide variety of different antigens (*11*). It is our definition that imagines the existence of an antigen (or pMHC) such that given an antibody-antigen pair (or TCR-pMHC pair), if a second antibody (or TCR) binds this antigen (or pMHC) half as well, the resulting similarity defines the relationship with the first antibody (or TCR) over all antigens (or pMHCs). Both basic biology (centered on the relationship between sequence and specificity) and experimental experience justify this line of reasoning. The conclusion can be illustrated by a counterfactual: it is not biologically reasonable that, as a general rule, two antibodies or TCRs bind all potential antigens or pMHCs wildly and/or unpredictably differently. Instead, each experimental measurement of relative binding for a pair of antibodies or TCRs is likely to be highly representative of the similarity landscape across all antigens or pMHCs for that pair (they both bind a given antigen or set of antigens fairly well, and both bind the millions of other, unrelated antigens for which they are not specific, at an orders-of-magnitude lower level, close to some baseline; this is illustrated by the low level of binding of the wide variety of negative controls across published ELISA studies). Populating the matrix ***Z*** using a similarity measure that averages over this landscape is left for future work.

A second limitation is imprecision in the prediction of similarity for a given pair of antibodies or TCRs. Despite great interest and progress in this area (*70, 71*), predicting ΔΔG, let alone measuring it at repertoire-scale throughput to populate similarity matrices (***Z***) with billions of entries, remains challenging. Fortunately, larger matrices benefit from the law of large numbers, making the repertoire-scale measures we report more reliable, and we found that small differences in class diversity between repertoires or over time can be reliably identified using conventional sample sizes. Larger public datasets of pairwise binding data would be beneficial. Finally, we note that the present study was limited to a single CDR, although the framework we describe is amenable to more comprehensive characterizations of antibodies and TCRs, or indeed any macromolecules, as such application requires simply updating the similarity matrix. Regardless, that the class diversity of an individual should so closely mirror the class diversity of a population, for both antibodies and TCRs, strongly supports the view that most individuals have similar antigen-binding capacity, and that the erstwhile “dark matter” of unshared or private genes organizes into public or shared binding classes. We expect that a better understanding of these classes, and of binding-based functional class overlap, will help characterize differences between individuals that may underlie differences in health or susceptibility to disease.

Our work highlights the need for larger training sets for predicting differences in binding. Despite SKEMPI being the best database available, its antibody and TCR data was insufficient basis for binding models based on specific amino-acid substitutions or biophysical properties. While such models fit training portions of the dataset better than *Z_ij_*=*s^m^*, they overfit this training data, resulting in same-or-worse predictions on the test portion of the training-test split. This is an example of why it is important to test models in this way, to reduce the risk of being falsely impressed by more “realistic” models whose additional realism or complexity are not in fact supported by data.

Overall, the results presented here illustrate the value and opportunities that can be unlocked by using repertoire-scale measures that are based on the defining function of repertoires’ elemental units: epitope binding, for both antibodies and TCRs. This approach was inspired by foundational and well-established ideas in immunology (*19, 34, 72*), ecology (*3, 32, 46, 73, 74*), and physics (*75, 76*). Class diversity differs from network-, lineage-, or cluster-based (*8, 65, 66*) descriptions of repertoires in that class diversity (*i*) avoids the need for similarity cutoffs, which are arbitrary but can have large effects on network architecture/cluster counts (*77*); (*ii*) accounts for weak antibody-antigen/TCR-pMHC interactions, which are the overwhelming majority and are considered important in immunology (*19-21*), as they are in other complex systems (*76*); and (*iii*) is based explicitly on a model of binding similarity (albeit a rough/limited one), as opposed to simply on nucleotide- or amino-acid edit distance (*65, 66*). Our method’s modular design means it can be easily updated using models of binding similarity (*Z_ij_*) that make use of additional sequence or structural data as such data becomes available, and is readily applied beyond immunology to measure, for example, class diversity of tumors (cell diversity) (*78*), microbiomes/metagenomes (bacterial/viral diversity) (*79*), and other complex systems (*80*). Classes redefine diversity.

## Author contributions

Both authors contributed to the conceptualization, data curation, formal analysis, funding acquisition, investigation, methodology, resources, software, validation, visualization, and writing. Dr. Arnaout provided supervision and funding acquisition.

## Acknowledgements

This work was supported by NIH NIAID R01AI148747; the Extreme Science and Engineering Discovery Environment (XSEDE), which is supported by NSF grant ACI-1548562; and the Comet supercomputer at the San Diego Supercomputer Center, which is supported by NSF grant ACI-1341698.

## Data and materials availability

All data needed to evaluate the conclusions in the paper are present in the paper, references, and/or the Supplementary Materials. Code is available upon request.

## Supplementary Materials

### Supplementary Figures

**Figure S1.**
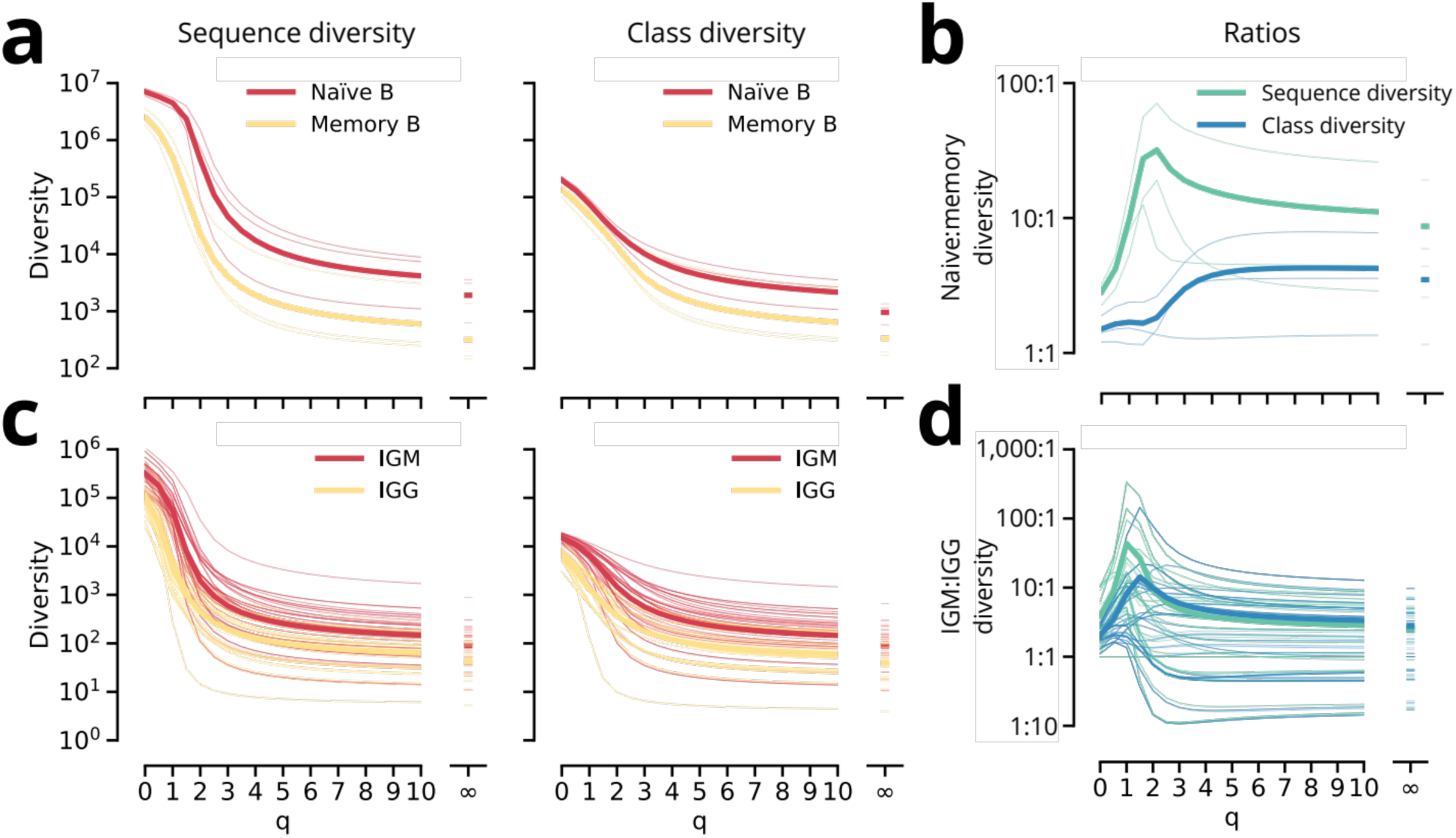
Sequence vs. class diversity of naive vs. memory and IGM vs. IGG repertoires. **(a)** Sequence vs. class diversity for exceptionally deeply sequenced naive vs. memory repertoires from DNA, and **(b)** the ratio of naive:memory diversity, as a function of *q* (see description of Equation 1 in the main text). Each thin line corresponds to a repertoire from a single individual. Each thick line is the average. **(c)-(d)** Analogous for IGM and IGG.

**Figure S2.**
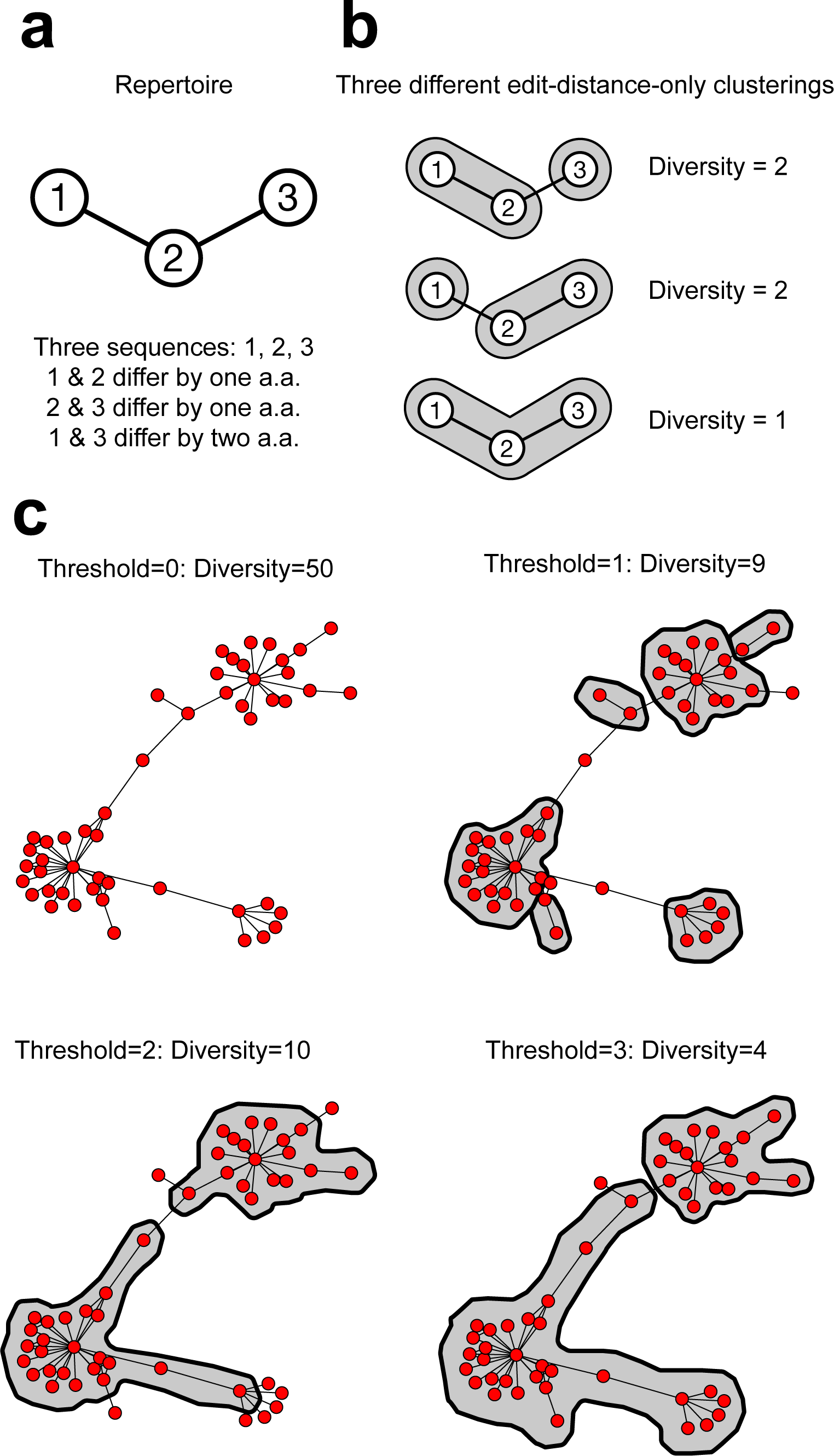
Class diversity ≠ edit distance: non-uniqueness of edit-distance-based diversity. We use a particular form of class diversity based on binding similarity; the similarity function we fit to the binding data in SKEMPI (*K_d_*) is what yields this particular form. However, *every* form of class similarity differs from edit distance in that class diversity is uniquely determined by its similarity function, whereas diversity measures based on edit distance alone—i.e., ones that are not based on a fit to any external data but are solely based on the number of clusters that result from a particular edit-distance cutoff—are not unique in this way. **(a)** shows the simplest “repertoire” that illustrates this point (Left). Each node represents a sequence. Edges connect sequences that differ at just a single amino-acid position. If we cluster by edit distance with a clustering threshold of 1 amino-acid difference, there are three different possible clusterings (right). In contrast, Eq. 1, which defines class diversity, gives a unique solution. **(b)** In edit-distance-only measures the clustering threshold need not be 1 amino acid; it can be 2, or 3, or indeed any arbitrary number. In contrast, the 0.3 in *Z_ij_*=0.3^m^ in the specific form of class diversity that we explore in this study is not chosen arbitrarily: it is the value determined by a fit to SKEMPI binding data. **(c)** Example of multiple different possible pure edit-distance-based diversity measures for a 50-sequence connected cluster from the Day 7 post-influenza-vaccination sample in Fig. S4. Each node is a unique sequence. Each pair of nodes is connected by an edge if they differ at a single amino-acid position. Here “diversity” means number of clusters at the indicated edit-distance threshold, beginning with the highest-degree node (the sequence with the most connections; same approach as in (b), middle panel). Clusters with more than 1 sequence are identified by a gray background. None of the shown thresholds convey that there are three related clusters. While some other edit-distance-based threshold or strategy could be used based on the topology, class diversity is not ad hoc or post hoc in this way, as it is based on independent data: binding data.

**Figure S3.**
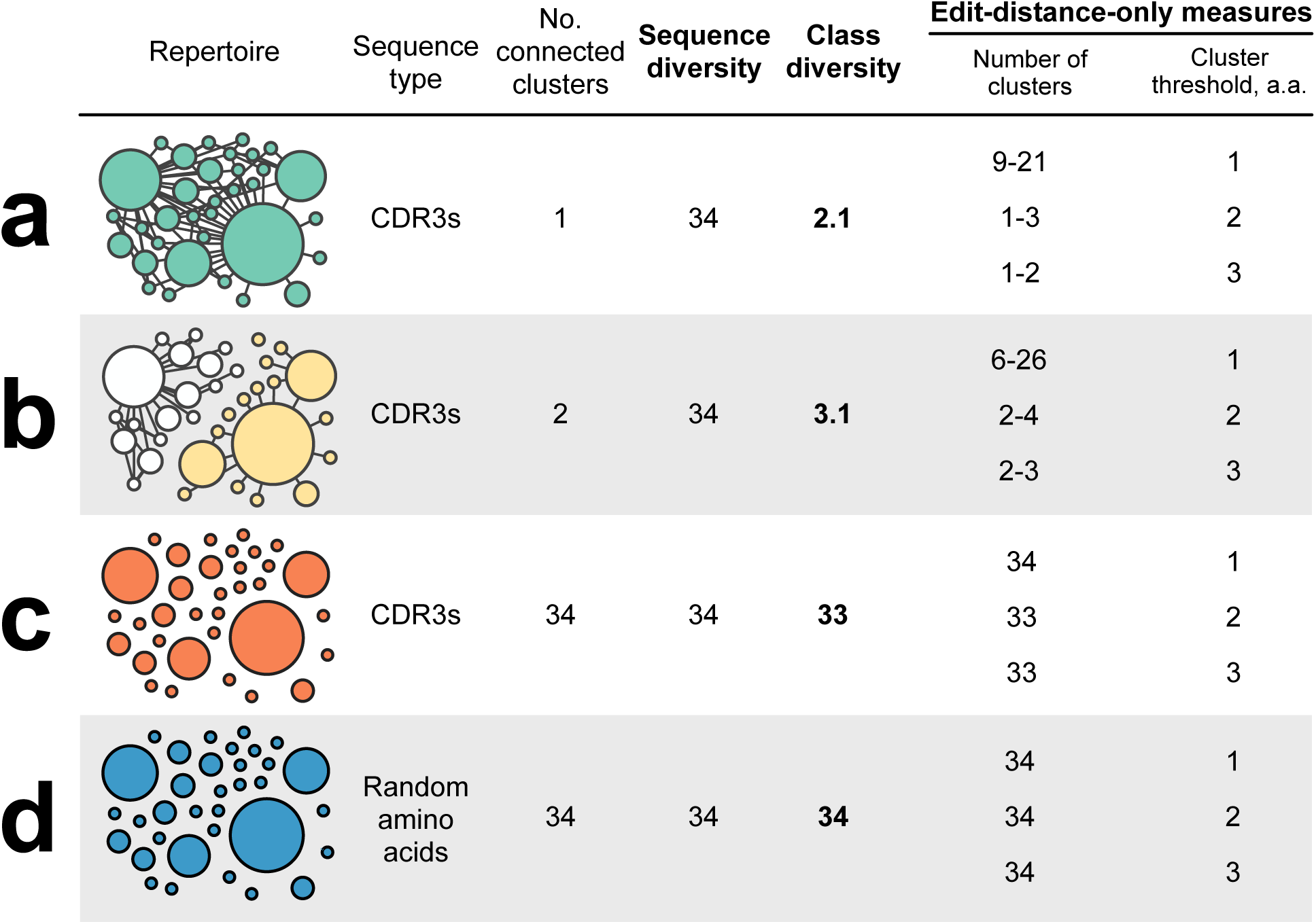
Class diversity vs. edit-distance thresholds. Edit-distance-based clustering requires a threshold be chosen: e.g., 1, 2, or 3 amino acids. Sequences that differ by this threshold amount or less are clustered together. The resulting number of clusters gives one measure of diversity. Different thresholds often give different clusters, and thereby different measures of diversity. This figure expands on Fig. 3d-g in the main text to illustrate how different cluster thresholds (last column) yield different numbers of clusters (second-to-last column) for the *in silico* repertoires from Fig. 3. Note the fairly wide ranges for repertoires **(a)** and **(b)**, a consequence of the non-uniqueness illustrated in Fig. S2. In the extremely diverse repertoires in **(c)** (all very different CDR3s) and **(d)** (random amino acids), edit distance approximates class diversity, but this happens only in the most extreme cases, not in typical repertoires (e.g. Fig. S4).

**Figure S4.**
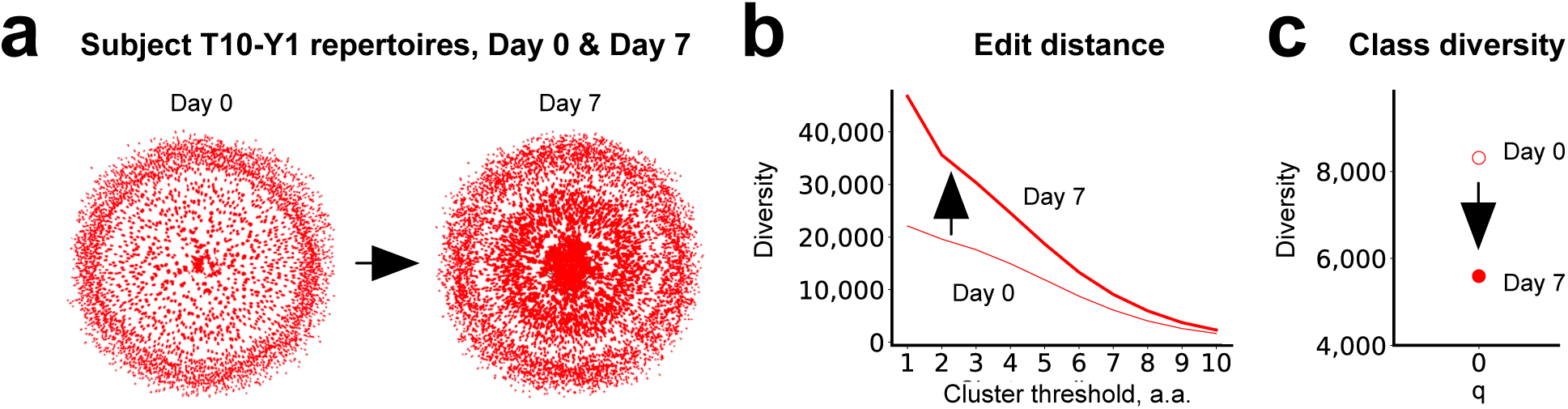
Binding-based class diversity does not correspond to a simple edit-distance threshold. **(a)** shows network representations of the day-0 and day-7 IGG CDR3 repertoires from subject T10-Y1 from Ref. 40: each dot represents a sequence, and each edge connects sequences that differ by a single amino-acid position. **(b)** Using purely edit distance, the diversity, measured as the number of clusters, ranges over orders of magnitude, depending on what cluster threshold is chosen. Absent more information, the choice of cluster threshold is arbitrary. Note that at every cluster threshold, diversity is higher in the Day 7 repertoire (thick line) than the Day 0 repertoire (thin line). **(c)** In contrast, there is no arbitrariness to the measures of class diversity presented in this study: they are determined by the fit to the *K_d_* binding data described. ^0^*D*S based on binding similarity happens to correspond roughly to a clustering threshold of 6 amino acids in the Day 0 repertoire and 8 amino acids in the Day 7 repertoire; there is no set correspondence because class diversity is different from simple edit-distance-based diversity (compare to thresholds of 1-3 amino acids in Fig. S3). Also in contrast to edit distance, class diversity in the Day 7 repertoire (filled symbol) is lower than in the Day 0 repertoire (open symbol), reflecting the capture of repertoire-wide structure that simple edit-distance-based measures cannot capture, at any clustering threshold.

## References

1. A. Six, M. E. Mariotti-Ferrandiz, W. Chaara, S. Magadan, H.-P. Pham, M.-P. Lefranc, T. Mora, V. Thomas-Vaslin, A. M. Walczak, P. Boudinot, The past, present, and future of immune repertoire biology - the rise of next-generation repertoire analysis. Front Immunol. 4, 413 (2013).

2. N. Chaudhary, D. R. Wesemann, Analyzing Immunoglobulin Repertoires. Front Immunol. 9 (2018), doi:10.3389/fimmu.2018.00462.

3. M. O. Hill, Diversity and Evenness: A Unifying Notation and Its Consequences. Ecology. 54, 427–432 (1973).

4. A. Hosoi, K. Takeda, K. Nagaoka, T. Iino, H. Matsushita, S. Ueha, S. Aoki, K. Matsushima, M. Kubo, T. Morikawa, K. Kitaura, R. Suzuki, K. Kakimi, Increased diversity with reduced “diversity evenness” of tumor infiltrating T-cells for the successful cancer immunotherapy. Scientific Reports. 8, 1–12 (2018).

5. O. V. Britanova, E. V. Putintseva, M. Shugay, E. M. Merzlyak, M. A. Turchaninova, D. B. Staroverov, D. A. Bolotin, S. Lukyanov, E. A. Bogdanova, I. Z. Mamedov, Y. B. Lebedev, D. M. Chudakov, Age-related decrease in TCR repertoire diversity measured with deep and normalized sequence profiling. J. Immunol. 192, 2689–2698 (2014).

6. J. J. Goronzy, C. M. Weyand, Mechanisms underlying T cell ageing. Nature Reviews Immunology. 19, 573–583 (2019).

7. S. D. Boyd, Y. Liu, C. Wang, V. Martin, D. K. Dunn-Walters, Human lymphocyte repertoires in ageing. Current Opinion in Immunology. 25, 511–515 (2013).

8. C. F. A. de Bourcy, C. J. L. Angel, C. Vollmers, C. L. Dekker, M. M. Davis, S. R. Quake, Phylogenetic analysis of the human antibody repertoire reveals quantitative signatures of immune senescence and aging. Proc Natl Acad Sci U S A. 114, 1105–1110 (2017).

9. D. Neri, S. Montigiani, P. M. Kirkham, Biophysical methods for the determination of antibody-antigen affinities. Trends Biotechnol. 14, 465–470 (1996).

10. A. C. Moser, S. Trenhaile, K. Frankenberg, Studies of antibody-antigen interactions by capillary electrophoresis: A review. Methods. 146, 66–75 (2018).

11. J. Jankauskaite, B. Jimenez-Garcia, J. Dapkunas, J. Fernandez-Recio, I. H. Moal, SKEMPI 2.0: An updated benchmark of changes in protein-protein binding energy, kinetics and thermodynamics upon mutation. Bioinformatics (2018), doi:10.1093/bioinformatics/bty635.

12. P. Dash, A. J. Fiore-Gartland, T. Hertz, G. C. Wang, S. Sharma, A. Souquette, J. C. Crawford, E. B. Clemens, T. H. O. Nguyen, K. Kedzierska, N. L. La Gruta, P. Bradley, P. G. Thomas, Quantifiable predictive features define epitope-specific T cell receptor repertoires. Nature. 547, 89–93 (2017).

13. J. Glanville, H. Huang, A. Nau, O. Hatton, L. E. Wagar, F. Rubelt, X. Ji, A. Han, S. M. Krams, C. Pettus, N. Haas, C. S. L. Arlehamn, A. Sette, S. D. Boyd, T. J. Scriba, O. M. Martinez, M. M. Davis, Identifying specificity groups in the T cell receptor repertoire. Nature. 547, 94–98 (2017).

14. J. A. Doering, S. Lee, K. Kristiansen, L. Evenseth, M. G. Barron, I. Sylte, C. A. LaLone, In silico site-directed mutagenesis informs species-specific predictions of chemical susceptibility derived from the Sequence Alignment to Predict Across Species Susceptibility (SeqAPASS) tool. Toxicol Sci. 166, 131–145 (2018).

15. I. Wirgin, N. K. Roy, M. Loftus, R. C. Chambers, D. G. Franks, M. E. Hahn, Mechanistic Basis of Resistance to PCBs in Atlantic Tomcod from the Hudson River. Science. 331, 1322–1325 (2011).

16. R. Albert, H. Jeong, A.-L. Barabási, Error and attack tolerance of complex networks. Nature. 406, 378–382 (2000).

17. K. W. Wucherpfennig, P. M. Allen, F. Celada, I. R. Cohen, R. De Boer, K. C. Garcia, B. Goldstein, R. Greenspan, D. Hafler, P. Hodgkin, E. S. Huseby, D. C. Krakauer, D. Nemazee, A. S. Perelson, C. Pinilla, R. K. Strong, E. E. Sercarz, Polyspecificity of T cell and B cell receptor recognition. Semin. Immunol. 19, 216–224 (2007).

18. M. Cohn, An in depth analysis of the concept of “polyspecificity” assumed to characterize TCR/BCR recognition. Immunol. Res. 40, 128–147 (2008).

19. A. Coutinho, M. D. Kazatchkine, S. Avrameas, Natural autoantibodies. Curr. Opin. Immunol. 7, 812–818 (1995).

20. M. Boes, Role of natural and immune IgM antibodies in immune responses. Mol. Immunol. 37, 1141–1149 (2000).

21. R. Reverberi, L. Reverberi, Factors affecting the antigen-antibody reaction. Blood Transfus. 5, 227–240 (2007).

22. P. L. Kastritis, A. M. J. J. Bonvin, On the binding affinity of macromolecular interactions: daring to ask why proteins interact. Journal of The Royal Society Interface. 10, 20120835 (2013).

23. T. Siebenmorgen, M. Zacharias, Computational prediction of protein–protein binding affinities. WIREs Computational Molecular Science. 10, e1448 (2020).

24. C. Geng, L. C. Xue, J. Roel-Touris, A. M. J. J. Bonvin, Finding the ΔΔG spot: Are predictors of binding affinity changes upon mutations in protein–protein interactions ready for it? WIREs Computational Molecular Science. 9, e1410 (2019).

25. E. D. Levy, A simple definition of structural regions in proteins and its use in analyzing interface evolution. J Mol Biol. 403, 660–70 (2010).

26. A. A. Bogan, K. S. Thorn, Anatomy of hot spots in protein interfaces. J Mol Biol. 280, 1–9 (1998).

27. H. M. Berman, J. Westbrook, Z. Feng, G. Gilliland, T. N. Bhat, H. Weissig, I. N. Shindyalov, P. E. Bourne, The Protein Data Bank. Nucleic Acids Res. 28, 235–242 (2000).

28. M. Sandberg, L. Eriksson, J. Jonsson, M. Sjöström, S. Wold, New chemical descriptors relevant for the design of biologically active peptides. A multivariate characterization of 87 amino acids. J. Med. Chem. 41, 2481–2491 (1998).

29. J. Dearden, The History and Development of Quantitative Structure-Activity Relationships (QSARs). International Journal of Quantitative Structure-Property Relationships. 1, 1–44 (2016).

30. R. Arora, J. Kaplinsky, A. Li, R. Arnaout, Repertoire-Based Diagnostics Using Statistical Biophysics. bioRxiv, 519108 (2019).

31. T. Leinster, C. A. Cobbold, Measuring diversity: the importance of species similarity. Ecology. 93, 477–489 (2012).

32. L. Jost, Entropy and diversity. Oikos. 113, 363–375 (2006).

33. C. Kahraman, B. Öztaysi, S. Ç. Onar, A Comprehensive Literature Review of 50 Years of Fuzzy Set Theory. Int. J. Comput. Intell. Syst. (2016), doi:10.1080/18756891.2016.1180817.

34. M. H. Van Regenmortel, Specificity, polyspecificity, and heterospecificity of antibody-antigen recognition. Journal of Molecular Recognition. 27, 627–639 (2014).

35. H.-P. Peng, K. H. Lee, J.-W. Jian, A.-S. Yang, Origins of specificity and affinity in antibody– protein interactions. PNAS. 111, E2656–E2665 (2014).

36. Q. Leng, Z. Bentwich, Beyond self and nonself: fuzzy recognition of the immune system. Scand. J. Immunol. 56, 224–232 (2002).

37. J. Kaplinsky, R. Arnaout, Robust estimates of overall immune-repertoire diversity from high-throughput measurements on samples. Nat Commun. 7, 11881 (2016).

38. A. Chao, Nonparametric estimation of the number of classes in a population. Scand. j. stat. 11, 265–270 (1984).

39. W. S. DeWitt, P. Lindau, T. M. Snyder, A. M. Sherwood, M. Vignali, C. S. Carlson, P. D. Greenberg, N. Duerkopp, R. O. Emerson, H. S. Robins, A Public Database of Memory and Naive B-Cell Receptor Sequences. PLoS ONE. 11, e0160853 (2016).

40. C. Vollmers, R. V. Sit, J. A. Weinstein, C. L. Dekker, S. R. Quake, Genetic measurement of memory B-cell recall using antibody repertoire sequencing. Proc. Natl. Acad. Sci. U.S.A. 110, 13463–13468 (2013).

41. A. L. Notkins, Polyreactivity of antibody molecules. Trends in Immunology. 25, 174–179 (2004).

42. S. Holmseth, Y. Zhou, V. V. Follin-Arbelet, K. P. Lehre, D. E. Bergles, N. C. Danbolt, Specificity Controls for Immunocytochemistry. J Histochem Cytochem. 60, 174–187 (2012).

43. Q. Chu, J. J. Ludtke, V. M. Subbotin, A. Blockhin, A. V. Sokoloff, The acquisition of narrow binding specificity by polyspecific natural IgM antibodies in a semi-physiological environment. Mol. Immunol. 45, 1501–1513 (2008).

44. G. J. Wedemayer, P. A. Patten, L. H. Wang, P. G. Schultz, R. C. Stevens, Structural Insights into the Evolution of an Antibody Combining Site. Science. 276, 1665–1669 (1997).

45. A. G. Schmidt, H. Xu, A. R. Khan, T. O’Donnell, S. Khurana, L. R. King, J. Manischewitz, H. Golding, P. Suphaphiphat, A. Carfi, E. C. Settembre, P. R. Dormitzer, T. B. Kepler, R. Zhang, M. A. Moody, B. F. Haynes, H. X. Liao, D. E. Shaw, S. C. Harrison, Preconfiguration of the antigen-binding site during affinity maturation of a broadly neutralizing influenza virus antibody. Proc Natl Acad Sci U S A. 110, 264–9 (2013).

46. L. Jost, What do we mean by diversity? The path towards quantification. Mètode Science Studies Journal - Annual Review. 0 (2018), doi:10.7203/metode.9.11472.

47. M. D. Cooper, M. N. Alder, The evolution of adaptive immune systems. Cell. 124, 815–822 (2006).

48. N. L. Bernasconi, E. Traggiai, A. Lanzavecchia, Maintenance of serological memory by polyclonal activation of human memory B cells. Science. 298, 2199–2202 (2002).

49. R. O. Emerson, W. S. DeWitt, M. Vignali, J. Gravley, J. K. Hu, E. J. Osborne, C. Desmarais, M. Klinger, C. S. Carlson, J. A. Hansen, M. Rieder, H. S. Robins, Immunosequencing identifies signatures of cytomegalovirus exposure history and HLA-mediated effects on the T cell repertoire. Nat Genet. 49, 659–665 (2017).

50. M. Ho, Epidemiology of Cytomegalovirus Infections. Rev Infect Dis. 12, S701–S710 (1990).

51. K. Peggs, S. Verfuerth, A. Pizzey, J. Ainsworth, P. Moss, S. Mackinnon, Characterization of human cytomegalovirus peptide-specific CD8(+) T-cell repertoire diversity following in vitro restimulation by antigen-pulsed dendritic cells. Blood. 99, 213–223 (2002).

52. W. H. Berger, F. L. Parker, Diversity of planktonic foraminifera in deep-sea sediments. Science. 168, 1345–1347 (1970).

53. K. Hashimoto, T. Kouno, T. Ikawa, N. Hayatsu, Y. Miyajima, H. Yabukami, T. Terooatea, T. Sasaki, T. Suzuki, M. Valentine, G. Pascarella, Y. Okazaki, H. Suzuki, J. W. Shin, A. Minoda, I. Taniuchi, H. Okano, Y. Arai, N. Hirose, P. Carninci, Single-cell transcriptomics reveals expansion of cytotoxic CD4 T cells in supercentenarians. PNAS. 116, 24242– 24251 (2019).

54. C. Soto, R. G. Bombardi, A. Branchizio, N. Kose, P. Matta, A. M. Sevy, R. S. Sinkovits, P. Gilchuk, J. A. Finn, J. E. Crowe, High frequency of shared clonotypes in human B cell receptor repertoires. Nature. 566, 398 (2019).

55. B. Briney, A. Inderbitzin, C. Joyce, D. R. Burton, Commonality despite exceptional diversity in the baseline human antibody repertoire. Nature. 566, 393–397 (2019).

56. M. Shugay, D. A. Bolotin, E. V. Putintseva, M. V. Pogorelyy, I. Z. Mamedov, D. M. Chudakov, Huge Overlap of Individual TCR Beta Repertoires. Front Immunol. 4 (2013), doi:10.3389/fimmu.2013.00466.

57. R. Cibotti, J. P. Cabaniols, C. Pannetier, C. Delarbre, I. Vergnon, J. M. Kanellopoulos, P. Kourilsky, Public and private V beta T cell receptor repertoires against hen egg white lysozyme (HEL) in nontransgenic versus HEL transgenic mice. J. Exp. Med. 180, 861–872 (1994).

58. G. Georgiou, G. C. Ippolito, J. Beausang, C. E. Busse, H. Wardemann, S. R. Quake, The promise and challenge of high-throughput sequencing of the antibody repertoire. Nat Biotechnol. 32, 158–68 (2014).

59. D. E. Bild, D. A. Bluemke, G. L. Burke, R. Detrano, A. V. Diez Roux, A. R. Folsom, P. Greenland, D. R. Jacob, R. Kronmal, K. Liu, J. C. Nelson, D. O’Leary, M. F. Saad, S. Shea, M. Szklo, R. P. Tracy, Multi-Ethnic Study of Atherosclerosis: objectives and design. Am. J. Epidemiol. 156, 871–881 (2002).

60. P. Parameswaran, Y. Liu, K. M. Roskin, K. K. Jackson, V. P. Dixit, J. Y. Lee, K. L. Artiles, S. Zompi, M. J. Vargas, B. B. Simen, B. Hanczaruk, K. R. McGowan, M. A. Tariq, N. Pourmand, D. Koller, A. Balmaseda, S. D. Boyd, E. Harris, A. Z. Fire, Convergent antibody signatures in human dengue. Cell Host Microbe. 13, 691–700 (2013).

61. J. Kaplinsky, A. Li, A. Sun, M. Coffre, S. B. Koralov, R. Arnaout, Antibody repertoire deep sequencing reveals antigen-independent selection in maturing B cells. Proc Natl Acad Sci U S A. 111, E2622–9 (2014).

62. R. Arora, R. Arnaout, Private Antibody Repertoires Are Public. BioRxiv, 2020.06.18.159699 (2020).

63. R. Arnaout, W. Lee, P. Cahill, T. Honan, T. Sparrow, M. Weiand, C. Nusbaum, K. Rajewsky, S. B. Koralov, High-resolution description of antibody heavy-chain repertoires in humans. PLoS One. 6, e22365 (2011).

64. R. A. Arnaout, Specificity and overlap in gene segment-defined antibody repertoires. BMC Genomics. 6, 148 (2005).

65. R. J. M. Bashford-Rogers, A. L. Palser, B. J. Huntly, R. Rance, G. S. Vassiliou, G. A. Follows, P. Kellam, Network properties derived from deep sequencing of human B-cell receptor repertoires delineate B-cell populations. Genome Res. 23, 1874–1884 (2013).

66. E. Miho, R. Roškar, V. Greiff, S. T. Reddy, Large-scale network analysis reveals the sequence space architecture of antibody repertoires. Nat Commun. 10, 1–11 (2019).

67. P. C. Tumeh, C. L. Harview, J. H. Yearley, I. P. Shintaku, E. J. Taylor, L. Robert, B. Chmielowski, M. Spasic, G. Henry, V. Ciobanu, A. N. West, M. Carmona, C. Kivork, E. Seja, G. Cherry, A. J. Gutierrez, T. R. Grogan, C. Mateus, G. Tomasic, J. A. Glaspy, R. O. Emerson, H. Robins, R. H. Pierce, D. A. Elashoff, C. Robert, A. Ribas, PD-1 blockade induces responses by inhibiting adaptive immune resistance. Nature. 515, 568–71 (2014).

68. L. G. Volkert, Enhancing evolvability with mutation buffering mediated through multiple weak interactions. BioSystems. 69, 127–142 (2003).

69. L. G. Volkert, M. Conrad, The Role of Weak Interactions in Biological Systems: the Dual Dynamics Model. Journal of Theoretical Biology. 193, 287–306 (1998).

70. M. Petukh, M. Li, E. Alexov, Predicting Binding Free Energy Change Caused by Point Mutations with Knowledge-Modified MM/PBSA Method. PLoS Comput Biol. 11, e1004276 (2015).

71. H. H. Lin, S. Ray, S. Tongchusak, E. L. Reinherz, V. Brusic, Evaluation of MHC class I peptide binding prediction servers: Applications for vaccine research. BMC Immunology. 9, 8 (2008).

72. A. S. Perelson, G. F. Oster, Theoretical studies of clonal selection: minimal antibody repertoire size and reliability of self-non-self discrimination. J. Theor. Biol. 81, 645–670 (1979).

73. A. Chao, C.-H. Chiu, L. Jost, Phylogenetic diversity measures based on Hill numbers. Philosophical Transactions of the Royal Society B: Biological Sciences. 365, 3599–3609 (2010).

74. S. M. Scheiner, A metric of biodiversity that integrates abundance, phylogeny, and function. Oikos. 121, 1191–1202 (2012).

75. Air Temperature (2020), (available at https://www.grc.nasa.gov/www/k-12/VirtualAero/BottleRocket/airplane/temptr.html).

76. M. S. Granovetter, The Strength of Weak Ties. American Journal of Sociology. 78, 1360– 1380 (1973).

77. V. Greiff, E. Miho, U. Menzel, S. T. Reddy, Bioinformatic and Statistical Analysis of Adaptive Immune Repertoires. Trends Immunol. 36, 738–749 (2015).

78. A. Heindl, C. Lan, D. N. Rodrigues, K. Koelble, Y. Yuan, Similarity and diversity of the tumor microenvironment in multiple metastases: critical implications for overall and progression-free survival of high-grade serous ovarian cancer. Oncotarget. 7, 71123– 71135 (2016).

79. K. Li, M. Bihan, S. Yooseph, B. A. Methé, Analyses of the Microbial Diversity across the Human Microbiome. PLOS ONE. 7, e32118 (2012).

80. R. Koopmans, M. Schaeffer, “De-composing diversity: In-group size and out-group entropy and their relationship to neighbourhood cohesion,” Discussion Papers, Research Unit: Migration, Integration, Transnationalization (WZB Berlin Social Science Center, 2013), (available at https://ideas.repec.org/p/zbw/wzbmit/spvi2013104.html).

81. X. Brochet, M. P. Lefranc, V. Giudicelli, IMGT/V-QUEST: the highly customized and integrated system for IG and TR standardized V-J and V-D-J sequence analysis. Nucleic Acids Res. 36, W503–8 (2008).

82. J. Dunbar, K. Krawczyk, J. Leem, C. Marks, J. Nowak, C. Regep, G. Georges, S. Kelm, B. Popovic, C. M. Deane, SAbPred: a structure-based antibody prediction server. Nucleic Acids Res. 44, W474–W478 (2016).

83. J. Leem, S. H. P. de Oliveira, K. Krawczyk, C. M. Deane, STCRDab: the structural T-cell receptor database. Nucleic Acids Res. 46, D406–D412 (2018).

84. J. Whittaker, A. V. Groth, D. C. Mynarcik, L. Pluzek, V. L. Gadsboll, L. J. Whittaker, Alanine scanning mutagenesis of a type 1 insulin-like growth factor receptor ligand binding site. J Biol Chem. 276, 43980–6 (2001).

85. Schrödinger, LLC, The PyMOL Molecular Graphics System, Version 2.0 (2015).

86. M. G. Taylor, A. Rajpal, J. F. Kirsch, Kinetic epitope mapping of the chicken lysozyme.HyHEL-10 Fab complex: delineation of docking trajectories. Protein Sci. 7, 1857–1867 (1998).

87. T. Scior, J. L. Medina-Franco, Q.-T. Do, K. Martínez-Mayorga, J. A. Yunes Rojas, P. Bernard, How to recognize and workaround pitfalls in QSAR studies: a critical review. Curr Med Chem. 16, 4297–4313 (2009).

88. H. Mei, Z. H. Liao, Y. Zhou, S. Z. Li, A new set of amino acid descriptors and its application in peptide QSARs. Biopolymers. 80, 775–786 (2005).

89. J. Tong, T. Che, Y. Li, P. Wang, X. Xu, Y. Chen, A descriptor of amino acids: SVRG and its application to peptide quantitative structure-activity relationship. SAR QSAR Environ Res. 22, 611–620 (2011).

90. C. C. Palliser, D. A. D. Parry, Quantitative comparison of the ability of hydropathy scales to recognize surface β-strands in proteins. Proteins: Structure, Function, and Bioinformatics. 42, 243–255 (2001).

91. H. Meirovitch, S. Rackovsky, H. A. Scheraga, Empirical Studies of Hydrophobicity. 1. Effect of Protein Size on the Hydrophobic Behavior of Amino Acids. Macromolecules. 13, 1398–1405 (1980).

92. K. W. Olsen, Internal residue criteria for predicting three-dimensional protein structures. Biochim Biophys Acta. 622, 259–267 (1980).

93. H. Cid, M. Bunster, E. Arriagada, M. Campos, Prediction of secondary structure of proteins by means of hydrophobicity profiles. FEBS Letters. 150, 247–254 (1982).

94. J. Kyte, R. F. Doolittle, A simple method for displaying the hydropathic character of a protein. J Mol Biol. 157, 105–132 (1982).

95. J. L. Cornette, K. B. Cease, H. Margalit, J. L. Spouge, J. A. Berzofsky, C. DeLisi, Hydrophobicity scales and computational techniques for detecting amphipathic structures in proteins. Journal of Molecular Biology. 195, 659–685 (1987).

96. H. Cid, M. Bunster, M. Canales, F. Gazitúa, Hydrophobicity and structural classes in proteins. *Protein Engineering*, Design and Selection. 5, 373–375 (1992).

97. P. K. Ponnuswamy, M. Prabhakaran, P. Manavalan, Hydrophobic packing and spatial arrangement of amino acid residues in globular proteins. Biochim Biophys Acta. 623, 301– 316 (1980).

98. Z. Botta-Dukát, The generalized replication principle and the partitioning of functional diversity into independent alpha and beta components. Ecography. 41, 40–50 (2018).

99. C.-H. Chiu, A. Chao, Distance-Based Functional Diversity Measures and Their Decomposition: A Framework Based on Hill Numbers. PLOS ONE. 9, e100014 (2014).

